# Investigating the occurrence of *E. coli* O157:H7 in the final effluents of two wastewater treatment plants

**DOI:** 10.1101/161737

**Authors:** Olayinka Osuolale, Anthony Okoh

**Affiliations:** Applied Environmental Metagenomics and Infectious Disease Research, (AEMIDR) Elizade University, Ilara-Mokin, Nigeria; SAMRC Microbial Water Quality Monitoring Centre, University of Fort Hare, Alice 5700, South Africa; Applied and Environmental Microbiology Research Group, Department of Biochemistry and Microbiology, University of Fort Hare, Alice 5700, South Africa

**Keywords:** E. coli O157:H7, Eastern Cape, Effluent, Wastewater, Latex Agglutination

## Abstract

**AIM:** The final effluent of two wastewater plants located in the Eastern Cape of South Africa were tested for the presence of Enterohaemorrhagic Escherichia coli O157:H7 (E. coli O157:H7) isolates, and characteristics of the isolates obtained were determined.

**METHODS AND RESULTS:** A total of 23 wastewater samples were collected from the treatment plants at the final effluent point after the disinfectant stages of wastewater processing. Altogether, 540 presumptive E. coli isolates were obtained by colony counting on the E. coli O157:H7 chromogenic agar base supplemented with cefixime tellurite and were sub-cultured onto sorbitol-MacConkey agar and tested for agglutination using the Prolex E. coli O157 latex test reagent kit. The results showed that the 149 suspected colonies from SMAC agar were all negative for the antisera.

**CONCLUSION:** None of the isolates agglutinated with antisera against E. coli O157. Thus no presence of the bacteria can be confirmed from the treated effluents

**SIGNIFICANCE AND IMPACT OF STUDY:** The likelihood of the receiving water body and the environment being contaminated with E. coli O157:H7 is therefore minimal.

## 1. Introduction

The most significant pathogenic *E. coli* for the water industry are the enterohaemorrhagic *E. coli* (EHEC). EHEC serotypes, such as *E. coli* O157:H7 and *E. coli* O111, produce large quantities of shiga-like (or vero) toxins that can cause diarrhoea that ranges from mild and non-bloody to highly bloody, which is indistinguishable from haemorrhagic colitis. Between 2% and 7% of cases can develop the potentially fatal haemolytic uraemic syndrome (HUS), which is characterized by acute renal failure and haemolytic anaemia. The infectivity of EHEC strains is substantially higher than that of other strains. As few as 100 EHEC organisms can cause infection ^1^, ^2^. EHEC comprise more than 100 different serotypes, including O157:H7, which has been responsible for a number of waterborne disease outbreaks ^2^. Current bacterial waterborne pathogens have been linked to gastrointestinal illnesses in human populations. Bacteria (e.g. *E. coli* O157:H7, Salmonella, Shigella, and Campylobacter jejuni), enteric viruses (e.g. Hepatitis A), and protozoa (e.g. Giardia, Cryptosporidium, Toxoplasmosis gondii) have all caused waterborne disease outbreaks^3^.

In the fall of 1991, Animal transmission, particularly from beef, had dominated O157:H7 outbreaks infections in South-Eastern Massachusetts provided an opportunity to identify transmission through beef as a seemingly unlikely vehicle ^4^. The *E. coli* O157:H7 was a culprit in a 2002 outbreak in Canada^5^. A more recent outbreak was a multi-state outbreak of the organism in November, 2013 in the US but the source remains under investigation ^6^. The first case of the *E. coli* O157 in South Africa was reported in the 1990s ^7^. Further attempts to isolate the organism in the wastewater treatment plant were performed in the Gauteng area of South Africa and gave low-positive confirmation ^8^. Muller, Ehlers and Grabow,^9^ reported infrequent incidence of *E. coli* O157 in the river water examined for direct and indirect domestic use in the Gauteng region of South Africa. In contrast, in the Eastern Cape of South Africa, Momba, Abong’o and Mwambakana,^10^ reported the isolation of *E. coli* O157:H7 in drinking water from 6 different communities.

The source of this organism has mostly been attributed to animals; cattle and possibly small domesticated ruminants constitute a primary animal reservoir ^11^. An outbreak in 1999 in Vancouver Washington traced the source of the E. coli O157:H7 to duck faeces ^12^. Ateba and Bezuidenhout,^13^ worked on livestock animals (cattle and swine faeces) and human faeces in South Africa, they reported higher percentage of the organism isolated and confirmed in the livestock faeces as compared to human faeces. The presence of the organism in human faeces was reported by Abong’o *et al.*,^14^ who isolated the organism from the stool of HIV/AIDS patients in the eastern cape of South Africa. A 3-year surveillance study revealed that in South Africa, less than 1% of the samples analysed, account for EHEC, concluding that EHEC is rarely isolated from humans in South Africa and is usually a coincidental finding ^15^. Abong’o,^16^ and Abong’o and Momba,^17^ reported the isolation of *E. coli* O157:H7 from meat and meat products in the eastern cape of South Africa.

Documentation of the incidence of infection from *E. coli* O157 in South Africa is scarce ^13^ and information on environmental detection of the organism is practically non-existing for South Africa in general and specifically for the province of study, the Eastern Cape. With these limited records and recent cases of outbreak of the *E. coli* in developed countries prompted the need to explore the possible presence of the organism in the Eastern Cape. Therefore, our aim was to evaluate the final effluent from the wastewater treatment plant as a means to identify occurrence of the organism in the environment.

## 2. Materials and Methods

### Wastewater treatment plant characteristics

The monitored wastewater treatment plants are located in the Buffalo City Municipality. Plant B sewage treatment works is situated in the Eastern Cape Province of the Buffalo City Municipality. It receives municipal domestic sewage as well as run-off water. The wastewater treatment plant operates an activated sludge system with design capacity of about 8ML/day which is considered a medium-sized treatment plant^18^. Presently, no industrial wastewater is discharged to the plant due to the closure of the industries in the area. The plant treats an average of dry weather flow of 7 000 m3/day and an average wet weather flow of 21 000 m3/day. In April 2013, an automated mechanical inlet Huber Screen was installed in the plant. This screen assists with the removal of rags and other foreign material from sewage before it enters the treatment process. There are two aeration tanks, each equipped with three vertically-mounted mechanical aerators, two anaerobic tanks and two clarifiers. A splitter box controls the flow of the raw sewage and return activated sludge (RAS) to the aeration tank. Sludge recycling is done through the RAS pump station hauling the sludge from the sedimentation tanks to the aeration tanks. The waste mixed liquid from the aeration tanks is pumped into the sludge lagoons. Chlorination is done by means of a water pressure operated, wall mounted, gas chlorinator in a baffle-resistant concrete contact tank. Thereafter, the final effluent is pumped to a pair of final effluent reservoirs ^19,20^ and discharged into a stream that links to the Mdizeni stream, tributary of the Keiskamma River ^18^.

Plant A is a medium size plant with treatment design capacity of 5Ml/d. The Bio-filter/Petroleum process treatment system is employed ^18^ and it discharge its final effluent in the Umzonyana stream.

### Sampling and analysis

Samples were collected on a monthly basis for 12 months from the final treated effluent (FE) and discharge point (DP). Samples were collected in one litre Nalgene bottles previously cleaned by washing in non-ionic detergent, rinsed with tap water, finally rinsed with deionised water and autoclaved prior to usage. Sodium thiosulphate was added to sampling bottles. Samples were then transported in cooler boxes containing ice packs to the Applied and Environmental Microbiology Research Group (AEMREG) laboratory at the University of Fort Hare, Alice, South Africa for analyses. Samples were processed within six hours of collection. Note, the sampling frequency and number of samples are as recommended by the Department of Water Affairs and Forestry ^21^. Sample blanks were used to analyse for any form of contamination serving as quality assurance. Sample blanks used were Trip blanks, Field blanks and Equipment blanks.

All blanks were analysed for the same parameters. All media (Selective media) used for bacterial analyses were performance-checked. Sterility and pH were checked as well as the proper reaction with appropriate positive and negative control organisms. Additionally, all bacterial tests were conducted with negative and positive controls run simultaneously with each assay. Test strains were bought from Leibniz-Institut DSMZ (GmBH).

Bacteriological analysis of the effluent samples for bacteria counts and isolation was determined by membrane filtration according to SABS^22^ and American Public Health Association^23^. Although turbidity of the effluent can severely inhibit the volume of water that can be passed through the filters, serial dilutions of the samples were prepared according to methods described by Rogers and Haines,^24^. In certain cases where there was excessive chlorine dosage in the effluent, the raw samples were filtered directly. Sample dilutions were homogenate before filtering. The filtered samples were placed on selective agar for the target organisms in triplicates. Petri dishes were allowed to dry for 15 minutes and inverted, and incubated promptly for 24hrs at 37°C. After 24 hrs incubation, counts in the suitable range (0-300 colonies) were recorded; using manual counting. The result per dilution plate counted was estimated.

### E. *coli* O157:H7 enumeration and isolation

*E. coli* O157:H7 was examined as described above. The filters were placed on *E. coli* O157:H7 chromogenic agar base (Conda, Madrid) supplemented with cefixime tellurite and incubated at 37°C for 24hrs. The target colonies appeared as pale pink in colour. They were counted and recorded as CFU/100 ml SABS^22^

### Preservation of Isolates

The isolates were taken as presumptive from the selective media on which they were grown base on their phenotypic identification and re-subculture for purification. Presumptive *E. coli* O157:H7 isolates was prepared in Luria broth and stored in 100% glycerol stock at -70°C.

### E. *coli* O157 latex agglutination assay

Each of the presumptive isolate was streaked onto a Sorbitol Mac-Conkey Agar supplemented with cefixime and tellurite plate from a 24hours growth culture and incubated for 18 h at 37°C. Sorbitol-negative or non-sorbitol fermenters (NSF) were tested for agglutination by the Prolex *E. coli* O157 latex test reagent kit (Pro-lab, Canada).

## 3. Results

Samples collected in the study were analysed for the presence *E. coli* O157:H7. Colonies with the characteristic pink/pale pink colour on the *E. coli* O157:H7 chromogenic agar were counted and taken as presumptive isolates for immunological testing. Overall, 41.7% of samples were positive for *E. coli* O157:H7 at Plant A treatment works and 45.8% were positive at the Plant B wastewater treatment plant. The bacterial counts varied from 1log10 to 3log10 for Plant B treatment plant and 1log10 to 4log10 for Plant A treatment works. The highest counts were observed in October and December, the period of rainfall (Data not shown). The months that were positive for the presumptive *E. coli* O157:H7 for each of the treatment plants are shown in Table 1.

**Table 1.** Showing the months positive and negative for the isolation of presumptive *E. coli* O157:H7.

| Site | Sep-12 | Oct-12 | Nov-12 | Dec-12 | Jan-13 | Feb-13 | Mar-13 | Apr-13 | May-13 | Jun-13 | Jul-13 | Aug-13 |
| --- | --- | --- | --- | --- | --- | --- | --- | --- | --- | --- | --- | --- |
| Plant A | ND<sup>1</sup> | + | + | + | - | - | - | - | - | + | + | - |
| Plant B | - | + | + | - | + | - | - | + | + | - | + | - |

<sup>1</sup> ND: not determined.

### 3.1. Latex agglutination testing

Of the 540 isolates (Plant A: 233, Plant B: 307) processed to determine the Non-Sorbitol fermenting (NSF) organisms using the CT-Sorbitol Mac-Conkey Agar, 149 (28%) yielded the characteristic NSF colonies. The NSF colonies were further tested using the latex agglutinating method. The latex agglutination assays were performed according to their manufacturers’ instructions. A positive result with the O157 latex reagents was interpreted as large clumps of agglutinated latex and bacteria with partial or complete clearing of the background latex within 1 to 2 min. All 149 NSF isolates were negative for the agglutination test (Table 2).

**Table 2.** Latex agglutination testing result for the presumptive isolates.

| Sampling site | Nature of sample | No of isolates tested | No (%) of isolates negative on Sorbitol Mac-Conkey (SMAC) | No of isolates positive for the latex agglutination test. |
| --- | --- | --- | --- | --- |
| Plant B | Effluent | 307 | 82(27%) | 0 |
| Plant A |  | 233 | 67(29%) | 0 |

## 4. Discussion

In the initial laboratory evaluation, the isolates tested positive for the *E. coli* O157:H7 chromogenic agar and the number reduced on CT-Sorbitol Mac-Conkey agar (CT-SMAC) and further testing of the NSF from the CT-SMAC was negative for the latex test. This ruled out the possibility of the presumptive isolates being *E. coli* O157:H7. Other literature has indicated that *Vibrio, Yersinia, Pseudomonas, Burkholderia* and *Aeromonas* spp. can also give the characteristic feature expected on the Sorbitol Mac-Conkey agar ^25,26^. Some other bacteria spp. have been reported to be non-sorbitol fermenters isolated from foods and stools ^27^. Hermos *et al.*,^28^ and Ngwa *et al.*,^29^ also found the media not to be selective enough for environmental samples. There are as well EHEC serotypes that are non-sorbitol fermenters (NSF) giving the characteristic growth on SMAC but are not H7 strain ^30^ well as non-O157 serotypes ^31^. While *E. coli* O157 are popularly known to be non-sorbitol fermenting organisms ^31^ there are reports of Sorbitol fermenting (SF) *E. coli* O157 with HUS ability from Czech^32^ and from Scotland ^33^.

In our region of study, the Eastern Cape, the presence of *E. coli* O157:H7 has been previously reported in drinking water. Drinking water analysed from Plant B was reported to have prevalence of the organism. Other areas of the Eastern Cape like Kwasaki, Fort Beaufort, Alice and Mdanstane were also reported to have *E. coli* O157:H7 in their drinking water. A molecular analysis of the presumptive isolates confirmed just a single isolate for Plant B, suggesting that there was no significant presence of the organism, *E. coli* O157:H7 ^16^.

Likewise, Muller, Clay and Grabow,^25^ tested for the presence of *E. coli* O157:H7 in the sewage plants and environmental samples from the Northern Province of South Africa and found the organism to be below the detectable limit. In the Gauteng Province of South Africa, Muller and Ehlers,^26^, detected low to no presence of the organism from both the sewage plants and environmental samples. Much of the reported cases of *E. coli* O157:H7 are attributed to food, dairy and animal products as major sources^34^. Work done in Turkey established cattle as the source for the prevalence of the *E. coli* O157:H7 as the organism was found in the slaughterhouse’s wastewater treatment plant^35^.

The environmental findings of this study, in which the occurrence of the organism in the final effluent could not be established, are in accordance with the low prevalence of *E. coli* O157:H7 infections among humans in the Southern Hemisphere, including South Africa, than in countries of the Northern Hemisphere ^26^. Other data on the occurrence and prevalence of EHEC are limited on outbreaks from water and environmental-related and their quantification or even the isolation has seldom been achieved by traditional or molecular methods^36^.

There is a great need for a more routine monitoring of wastewater treatment plants expanding the investigation to the Sorbitol fermenting EHEC as this study only focused on the non-sorbitol fermenting EHEC. Also studies on the dairy farms in the Eastern Cape provide an avenue to study the presence of the NSF, SF EHEC and non-O157 and their impact on the surrounding environmental resources. This monitoring needs to be done in terms of EHEC in the Eastern Cape of South Africa and how to safeguard public health.

In this work, the occurrence of the *E. coli* O157:H7 in the final effluent of the two wastewater plants could not be found. With the variability in the selective media used, it is possible that other *E. coli* O157 colonies were missed which included the NSF O157 as again the SF O157 and non-O157, that may be high in prevalence than none that was seen in the two different final effluent samples. On the other hand, it can also be possible that the prevalence of *E. coli* O157:H7 in the samples investigated in this study was indeed low. Lack of information and literature on the possible presence of these pathogens from wastewater treatment plants especially in the Eastern Cape makes it impossible to ascertain the potential presence of the organism. There are no reported cases of clinical infections caused by the organism in this region.

## Acknowledgments

The authors would like to thank the Water Research Commission of South Africa (Grant K5/2145), and the South African Medical Research Council for financial support.

## Author Contributions

Olayinka Osuolale conducted all sampling and experiments and wrote the manuscript; and Anthony Okoh supervised the project and corrected and edited the manuscript.

